# Optogenetic activation of muscle contraction *in vivo*

**DOI:** 10.1101/2020.07.07.192377

**Authors:** Elahe Ganji, C. Savio Chan, Christopher W. Ward, Megan L. Killian

## Abstract

Optogenetics is an emerging alternative to traditional electrical stimulation to initiate action potentials in activatable cells both ex vivo and in vivo. Optogenetics has been commonly used in mammalian neurons and more recently, it has been adapted for activation of cardiomyocytes and skeletal muscle. Therefore, the aim of this study was to evaluate the stimulation feasibility and sustain isometric muscle contraction and limit decay for an extended period of time (1s), using non-invasive transdermal light activation of skeletal muscle (triceps surae) in vivo. We used inducible Cre recombination to target expression of Channelrhodopsin-2 (ChR2(H134R)-EYFP) in skeletal muscle (Acta1-Cre) in mice. Fluorescent imaging confirmed that ChR2 expression is localized in skeletal muscle and does not have specific expression in sciatic nerve branch, therefore, allowing for non-nerve mediated optical stimulation of skeletal muscle. We induced muscle contraction using transdermal exposure to blue light and selected 10Hz stimulation after controlled optimization experiments to sustain prolonged muscle contraction. Increasing the stimulation frequency from 10Hz to 40Hz increased the muscle contraction decay during prolonged 1s stimulation, highlighting frequency dependency and importance of membrane repolarization for effective light activation. Finally, we showed that optimized pulsed optogenetic stimulation of 10 Hz resulted in comparable ankle torque and contractile functionality to that of electrical stimulation. Our results demonstrate the feasibility and repeatability of non-invasive optogenetic stimulation of muscle in vivo and highlight optogenetic stimulation as a powerful tool for non-invasive in vivo direct activation of skeletal muscle.

## Introduction

Contactless optical control of action potentials (APs) using optogenetics is an emerging alternative to traditional electrical stimulation for initiating contractions in skeletal and smooth muscles. Optogenetics utilizes light responsive microbial photosensitive proteins (opsins) to translocate ions across lipid membranes, enabling precise (targeted cell-specific) and fast (millisecond [ms]-time scales) optical control of membrane voltages.^1,2^ Channelrhodopsin-2 (ChR2) is a commonly used seven-layer transmembrane example of the light-sensitive opsins that acts as a non-specific inward cation channel (depolarizing) with an intrinsic light sensitivity.^3,4^ Targeted expression of ChR2 has been commonly used as a tool for the repeatable and high-kinetic targeted activation of, most commonly, mammalian neurons^1,2,5^ and, more recently, cardiomyocytes^6^ with light. In heart, ChR2 expression in cardiomyocytes has been mediated by non-viral delivery of exogenous light-sensitive ChR2 (e.g., tandem cell units)^7^; viral ChR2 expression with CAG promoters in embryonic stem cells^8^; and transgenic mouse lines expressing ChR2.^8^ Enabled by these approaches, optical activation has been shown a promising strategy to address cardiac arrhythmia and explore cardiac development.^8,9^ Optogenetics applications, Typically, muscle activation is initiated by the motor neuron whose voltage-dependent release of acetylcholine into the neuromuscular junction (NMJ) initiates the sodium (Na^+^) channel activation and local post-synaptic cation influx (Na^+^) and efflux (K^+^) the muscle cell transmembrane whose threshold depolarization initiates action potentials (APs) in the muscle fiber, however, are not limited to neurons and cardiac muscle cells.

Optogenetics is a promising tool in skeletal muscle activation both ex vivo and in vivo. Skeletal muscle activation can be achieved approaches either using a neuron-mediated excitation through presynaptic ion channels (indirect; electrical and optogenetic stimulation) or transmembrane-mediate excitation through post-synaptic channels (direct; optogenetic stimulation). Ex vivo, synchronous contraction at high (tetanus-like, sustained contraction) and low frequency (twitch-like contraction) has been induced by photoactivation of ChR2 expressing multinucleated myotubes (C2C12).^10^ Contactless optogenetic control of myotube contraction, with high temporal resolution, circumvents technical challenges faced in ex vivo electrical stimulation (e.g. electrode electrolysis and tissue damage).^10^ *In vitro*, contraction in explanted soleus muscle is induced with photoactivation of mouse skeletal muscle with targeted expression of ChR2 in enabled of muscles *in vitro*.^11^ *In vivo*, light-stimulated control of mice with non-specific expression of ChR2 in skeletal muscle sarcolemma and T-tubule (Sim1-Cre) has been studied for restoration of function in denervated muscles.^12^ Electrical stimulation, although with high temporal resolution in neural stimulation, lacks the spatial specificity of optogenetic stimulation in targeting cell subpopulations based on morphological markers and molecular footprint even in single cell intracellular electrical-stimulation approaches, are limited to in vitro scenarios and cannot be used simultaneous in vitro stimulation of multiple single neurons or *in vivo*.^13,14^ Optogenetics, on the contrary has excellent spatiotemporal resolution and is cell-specific light-responsiveness to change transmembrane voltage.^1,2^ This spatiotemporal specificity of optogenetics, prompts its potential to be utilized in conjunction with denervation models, using either indirect approaches that target secondary motor cortex or sub-population of neurons in the peripheral nervous system^15^ or using direct activation of muscle cell, sarcolemma, or T-tubules.^12^ Finally, optogenetic activation of skeletal muscle has potential in mitigating some of these existing limitations in electrical stimulation approaches (i.e. tissue-damage caused by repeated needle electrode placement, especially in juvenile animals, and limited applicability in conjunction with denervation models).

Herein we report our development and validation of an optogenetic method for the direct (non-nerve mediated) activation of skeletal muscle *in vivo*. The goal of this study was to develop a method for sustaining muscle contraction using non-invasive approaches (e.g., optogenetics) and limit decay in force generation for an extended period. We used Cre recombinase to induce targeted expression of ChR2(H134R)-EYFP in Acta1-Cre mice and an optimized experimental setup for *in vivo* muscle illumination in order to sustain contraction at the supraphysiological activation duration of 1 second, and we showed the feasibility and repeatability of targeted activation of the skeletal muscle *in vivo*.

## Methods

### Animal model

All procedures were approved by the Institutional Animal Care and Use Committee at the University of Delaware (N = 21 mice). Mice were housed in same-sex cages (maximum of five per cage) with littermates in BSL1 containment and monitored daily with regular chow and water *ad libitum* with a 12hr on/12hr off light cycle. We used *in-vivo* Cre-lox recombination for targeted expression of channelrhodopsin-2/YFP (ChR2(H134R)-EYFP) fusion protein (Ai32) in skeletal muscle.^16^ To generate these mice, we crossed doxycycline-inducible, skeletal muscle-specific ACTA1-rtTA,tetO-cre male mice (Acta1-Cre; C57BL6J background) with Ai32 reporter female mice (C57BL6J background) (Figure 1A).^17,18^ Doxycycline chow (Envigo) was provided *ad libitum* at time of mating to dams and continued until pups were weaned at 3-4 weeks of age. Offspring were genotyped using PCR (Transnetyx, TN, USA). Acta1-Cre; Ai32 homozygous mice (experimental) were used for blue light-induced stimulation and heterozygous or homozygous Ai32 (Cre-negative) offspring were used as controls. To verify Acta1-Cre specificity to skeletal muscle, we similarly crossed Ai14 (tdTomato) reporter mice with Acta1-Cre mice to generate Acta1-Cre; Ai14 reporters (N = 2).

**Figure 1:**
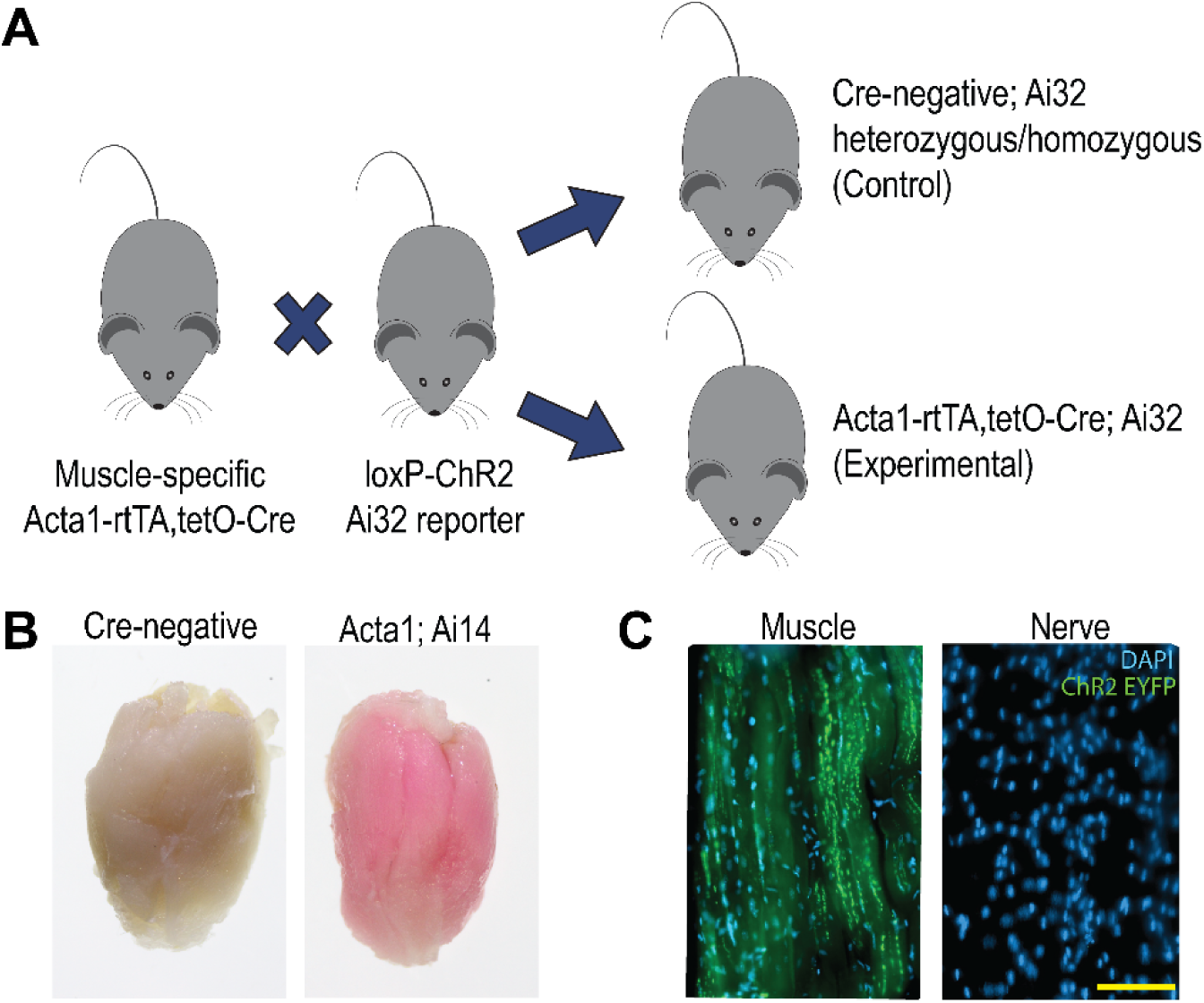
(A) Schematic showing the breeding scheme used for generation of experimental (Acta1-Cre Ai32) and control (Cre-negative) mice. (B) Whole-mount triceps surae muscle (TS) of Control and Acta1-Cre; tdTomato mice. (C) ChR2 was expressed in the skeletal muscle fibers but was not expressed in the sciatic nerve using the Acta1-promoter. Scale bar denotes 100 μm.

### ChR2 location and nerve infiltration

To visualize localization of the ChR2(H134R)-EYFP protein in the skeletal muscle and nerve, we dissected the quadriceps muscle and the main femoral branch of the sciatic nerve from Acta1-Cre; Ai32 mice at euthanasia (N = 2). Harvested tissues where fixed in 4% PFA, cryoprotected in sucrose, mounted in OCT freezing medium (Sakura, CA, USA), and sectioned at 30 µm thickness. Collected slides were then imaged with an Imager A2 microscope (Carl Zeiss, Germany).

### Optogenetic stimulation setup

We constructed a custom system with a LED (455 nm, 900 mW, M455L3, Thorlabs, NJ, USA), a collimator (SM2F32-A; 350-700 nm, Thorlabs, NJ, USA) to reduce energy dissipation, and high-power 1-Channel LED Driver (DC2200, Thorlabs, NJ, USA) for pulse modulation. The LED driver interface was custom programmed to induce user-defined pulsatile light activation intervals using LabView (National Instruments, TX, USA).

### Mouse preparation

Mice were anesthetized using isoflurane administration (1-2%). Hair was removed from hindlimb at the triceps surae muscle using chemical hair remover (Nair, Church & Dwight Co., NJ, USA). Animals were placed on a heating pad (Stoelting, IL, USA) at 37 °C in the prone position and their hindlimb was placed in a modular *in-vivo* muscle stimulation apparatus (Aurora Scientific, ON, Canada). The foot was placed in the footplate connected to a servomotor, with the ankle axis of rotation aligned with and tibial axis perpendicular to the servomotor shaft.

The hindlimb was then clamped at the knee to stabilize the joint construct at 90° tibiofemoral joint angle for isometric muscle contractions. Real-time measurements of applied ankle torque to the foot pedal during light-induced plantar flexion were collected with a 1 N force transducer (Aurora Scientific, ON, Canada).

### Muscle testing paradigm

We used an iterative process to determine suggested optogenetic stimulation parameters to increase the light-induced generated peak isometric ankle torque and obtain sustained submaximal contractions. Short-duration (twitch-like) light exposure was administered to exposed skin. The ankle torque was measured and the distance between skin and LED and the collimator lens displacement were selected at the corresponding values to higher measured isometric ankle torque readouts. Subsequently, the stimulation protocol (on and off) was adjusted to increase peak isometric ankle torque readout. To determine the frequency for sustained muscle contraction and minimize decay in contraction profile, we evaluated peak isometric torque at each stimulation frequency and ankle torque profiles in male Acta1-Cre; Ai32 mice (N = 3; Male; 3 months of age). The triceps surae muscles were stimulated at two different frequencies of 10 Hz (70 ms on-/30 ms off-time; 10 repetitions; 1 second total duration) and 40 Hz (20 ms on-/5 ms off-time; 40 repetitions; 1 second total duration). We selected 10 Hz stimulation as the optimized stimulation protocol for sustaining muscle contraction and used this protocol for the remainder of this study.

### Optimization and repeatability experiments

Variations in optogenetic stimulation are dependent on: (1) strength of stimulus (power of LED), (2) duration of stimulus (frequency used in protocol), and (3) cell type^17^. Therefore, we evaluated the efficiency of our optogenetic setup by (1) measuring penetration of light (power intensity) through the skin with the 10 Hz stimulation protocol; (2) measuring weight-dependent generation of ankle torque, (3) assessing repeatability using contralateral limb experiments, and (4) assessing contractile function of 10 Hz optogenetic stimulation compared to a comparable duration of electrical stimulation.

### Power meter experiments

We quantified the transmission of emission light intensity with a thermopile powermeter sensor (Coherent PM10; 19 mm), with an appropriate wavelength sensitivity to the LED of use (power 1150-1145 mW) to a and power meter (Fieldmate, Coherent). When placed at the experimental fixed distance from the light source (Figure 2B), the percentage of light intensity loss through the skin was determined through fresh dissected hairless skin patches from the abdomen and back of euthanized mice. In each case, power readouts divided by active sensor area (19 mm) to calculate light intensity.

**Figure 2:**
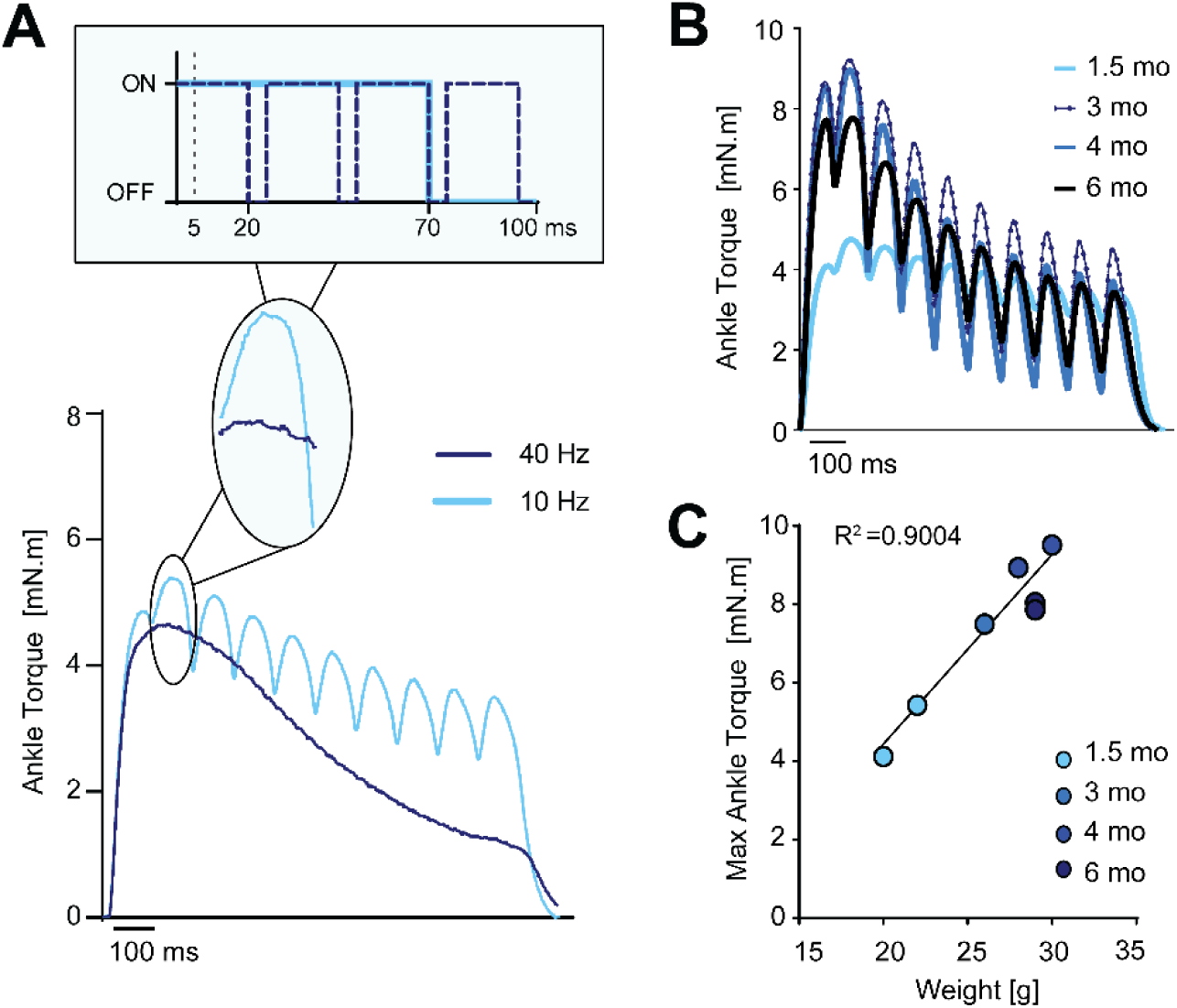
(A) Representative ankle torque profiles during 1 sec duration stimulation, comparing 10 Hz and 40 Hz. (B) Representative profile of generated ankle torque using 70 ms on/30 ms off blue light stimulation for different aged mice. (C) Light-activated ankle torque was linearly correlated with mouse weight (R^2^=0.9004).

### Weight-dependent response of optogenetic activation

The weight-dependence of optogenetic stimulation *in vivo* was assessed in Triceps surae muscle of Acta1-Cre; Ai32 male mice at 1.5, 3, 4, and 6-months of age (N = 2/age). Optical stimulation was at 10 Hz protocol as previously described with ankle torque recorded.

### Electrical vs. optogenetic activation

The right hindlimb of adult Acta1-Cre; Ai32 mice (N = 4; 2 males and 2 females, >3-months of age) was randomly assigned to either optogenetic or electrical activation of the Triceps surae with the contralateral limb assigned to the other. Optogenetic stimulation was a 10 Hz stimulation protocol (70 ms on time, 30 ms off time, 5 reps), electrical stimulation was with a tetanic pulse trains (0.3 (ms) pulse width, 150 (Hz) pulse frequency, 0.5 (s) duration) via sub-dermal PTFE-coated stainless steel EMG needle electrodes placed at the posterior knee and distal Achilles to activate the tibial nerve. We chose not to sever peroneal nerve for inhibition of antagonist anterior crural muscles contraction to maintain a non-invasive stimulation protocol comparable the stimulation using optogenetics. Stimulations were repeated for three consecutive days with ankle torque as the end-point measure. Measured torque values (mN.m) were normalized to body mass (kg) to account for mouse weight variability.

### Statistical analysis

Peak measured ankle torque values from electrical and optogenetic stimulation were statistically analyzed using two-way ANOVA with repeated measures or paired t-tests. The relationship between bodyweight and peak measured ankle torque was determined by linear regression. All statistical analyses were performed in Prism (version 7, Graphpad, LaJolla, CA).

## Results

### Confirmation of Acta1-Cre specificity and expression of Ai32 in muscle

Acta1-Cre Ai14 mice expressed robust tdTomato fluorescence in skeletal muscle following cre recombination, verifying the lineage tracing of Acta1-Cre (Figure 1B). Results from fluorescence imaging of nerve and skeletal muscle also showed that while ChR2(H134R)-EYFP was strongly expressed in skeletal muscle, it had no specific expression in the main femoral sciatic nerve branch (Figure 1C).

### Optimization

We were able to induce muscle contraction using transdermal exposure to blue light with Light On-Off optimization experiments (Supplemental Figure 2A, B) revealing 70 ms-On:30 ms -Off as optimal. Furthermore, we show that 10 Hz light stimulation for 1000 ms elicited a partially fused yet sustained tetanic plateau with higher frequency (40 Hz) exhibiting tetanic plateau decay (Figure 2A). Despite variable plateau decay, peak force was stable during repeated optogenetic stimulation (Supplemental Figure 3C). Despite a 74-80% reduction of light intensity through abdominal skin and 91% reduction through back skin, we showed that the light intensity was maintained at 0.26 and 0.35 mW/mm^2^ respectively (Figure 3).

**Figure 3.**
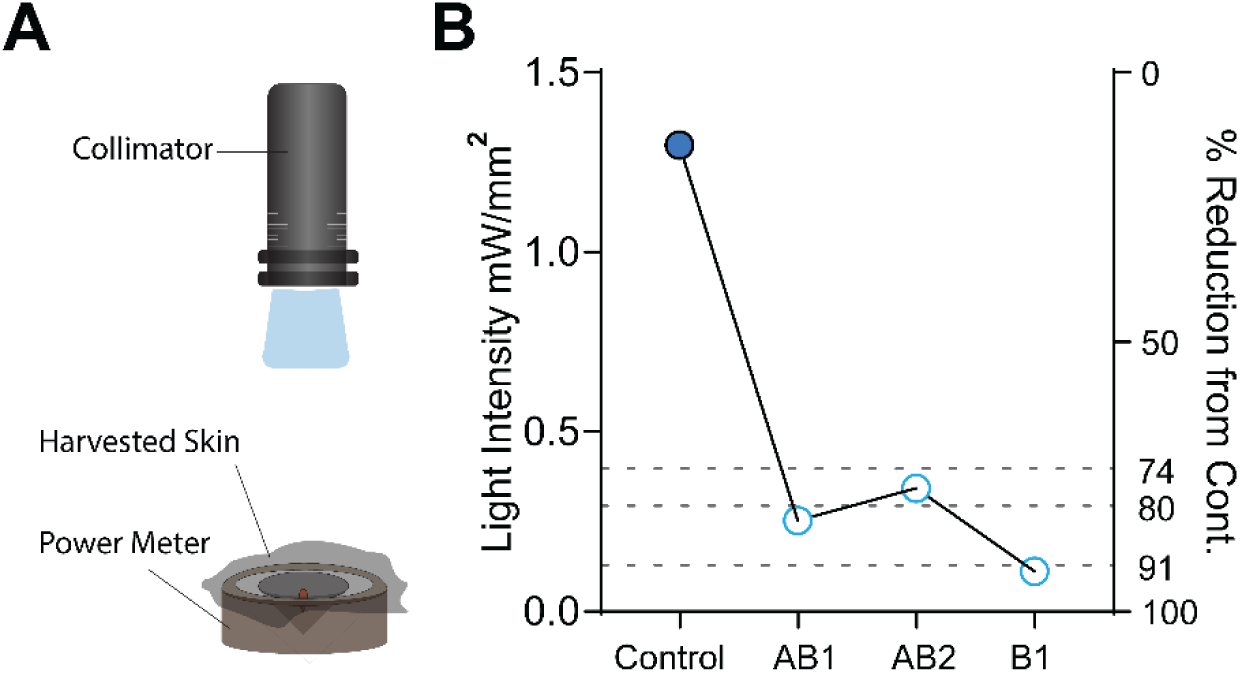
(A) Schematic of the experimental setup for light intensity measurements through collected skin. (B) Although the transdermal delivery of blue light reduced the light intensity, it still maintains an intensity of 0.2-0.3mW/mm^2^. AB = abdominal skin; B = back skin.

The magnitude of generated ankle torque under blue light stimulation correlated with the weight of the mice, as expected (Figure 2B and C, R^2^ = 0.9004, p < 0.001). Our method also generated repeatable muscle contractions in contralateral limbs (Supplemental Figure 3A, B) and in young adult mice of differing ages.

Finally, in comparison to electrical stimulation of tibial nerve (Figure 4A), our optimized pulsed optogenetic stimulation of 10 Hz (Figure 4B) resulted in comparable ankle torque values (Figure 4C) and measurements are repeatable for each stimulation protocol across three consecutive days.

**Figure 4.**
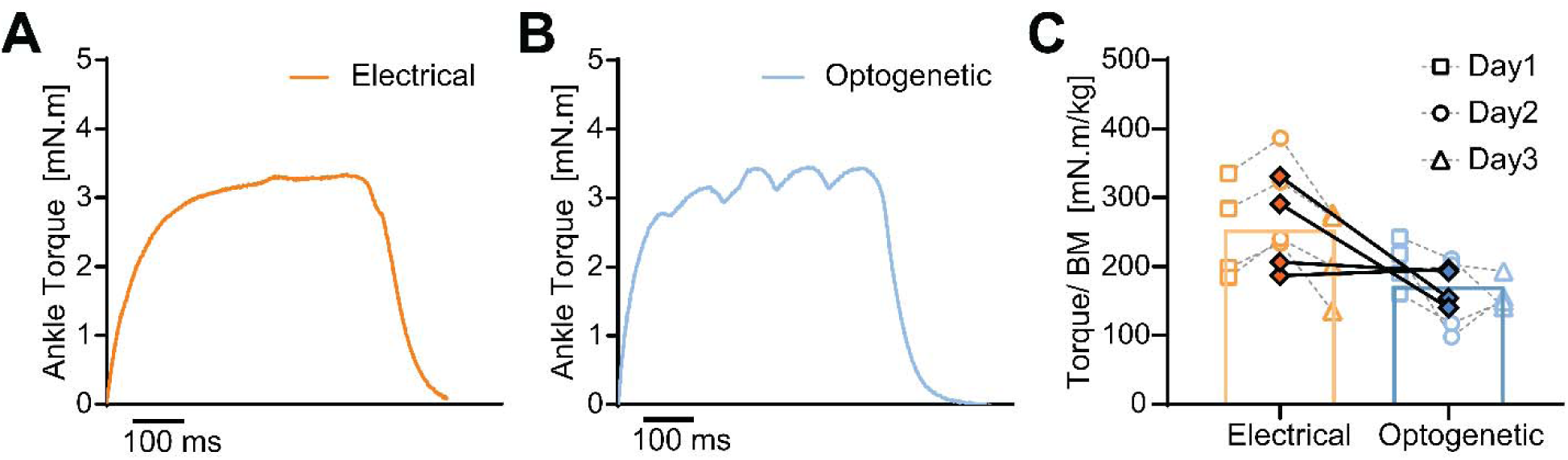
(A) Representative ankle torque profiles for electrical stimulation (0.3 ms pulse duration; 150Hz frequency; 0.5 sec duration) and (B) optogenetic stimulation (0.5 sec duration; 10Hz). (C) The average peak ankle torque generated during stimulations was comparable or slightly reduced for optogenetics stimulation compared to electrical stimulation.

## Discussion

Our findings demonstrate the feasibility and repeatability of optogenetics for non-invasive plasmalemmal stimulation of skeletal muscle using *in vivo*. Using cre-lox recombinase system, we induce targeted expression of channelrhodopsin-2/YFP in skeletal muscle (Acta1-Cre). We showed that targeted expression of ChR2 in Acta1-Cre enables nerve-independent activation of skeletal muscle *in vivo* and has comparable contractile function to that induced by electrical stimulation of tibial nerve in native muscle construct.

We confirmed the feasibility of muscle contraction via direct light exposure through the skin, despite light intensity reduction, and measured the light intensity. Light transmission through hairless abdominal and back skin maintained an intensity of 0.2-0.3mW/mm^2^ that is in agreement with the previously reported minimum light intensity of 0.35 mW/mm^2^ in skeletal muscle stimulation for soleus muscle explants without first passing light through skin.^11^ In this study, we showed that our experimental setup exceeds the minimum delivery of light intensity from our blue LED light source even after passing through skin from the trunk, which is more thick than skin from the limbs.

The goal of our experiments was to develop a method for sustaining muscle contraction using non-invasive approaches (e.g., optogenetics) and limit decay in force generation for an extended period of time. We successfully optimized our experimental setup by controlling the light intensity and pulse duration because we found that continuous illumination of light resulted an unsustainable contraction force profile. As we increased pulse duration, we found that the maximum force generation increased until we reached 70ms duration, which matched previously-reported findings by Bruegmann et al.^11^ In this same study, Bruegmann et al. also linked the repetition rate of light stimulation to effective depolarization and repolarization of membrane potentials in order to generate sustained muscle contraction, where repetition rates >40 Hz reduced the efficiency of optogenetic stimulation.^11^ The duty cycle of on/off time for optogenetic stimulation is a critical factor for generating muscle contraction. Here, we showed that the duration of light exposure generated the largest relative ankle torque at ∼70ms, and this finding was similar to previously reported durations of >70ms for ChR2 receptors to induce depolarization more efficiently (at lower threshold) compared to electrical stimulation in cardiomyocytes.^19^ The off-duty cycle is also important, as it allows the membrane to repolarize. In this study, we found the decay profile to be sustained at higher forces when using a longer (30ms) off-time between individual light pulses compared to shorter (10ms) off-time. This finding aligned with work by others that reported a deactivation time constant for ChR2 at ∼20 ms in order to obtain sufficient repolarization of lipid membrane between activations.^12^ The combination of both on- and off-duty cycling times are likely synergistic, and our comparisons of 10Hz and 40Hz repetition rates underscores the importance of repolarization for maintaining sustained contraction. Based on results from our study and two previous studies of optogenetic activation in intact mammalian skeletal muscle, we suggest that an effective illumination protocol (including both light intensity and pulsatile stimulation parameters) must be found based on anatomical site, ChR2 expression method, and experimental approach in future studies.^11,12^

Finally, in this study, we showed that using an optimized optogenetic stimulation protocol (10 Hz), we can induce increased and sustained muscle force generation, comparable to that of electrical stimulation. It is noteworthy that the results from electrical stimulation of the hindlimb in our study are lower in comparison to reported values in literature in stimulation of plantar flexor muscle construct with peroneal nerve severed to inhibit contraction.^20^ However, this may be explained by antagonist muscle contraction because, in our study, we kept the peroneal nerve intact during our hind limb stimulation. We made this choice in order to keep the *in vivo* muscle complex intact and comparable to optogenetic stimulation setup.

Challenges in maximal and repeatable direct optogenetic stimulation are linked to: (1) restricted recruitment of muscle fibers due to light absorption of myoglobin^11^ and (2) differences in myotube maturation level and the response of myotubes to excitation.^10^ Responsive cell distribution and ChR2 expression level are fundamental for efficiency of opsin excitation, as overexpression of ChR2 and its variants are needed to overcome low conductance and desensitization of the opsin. Yet high opsin expression can be disruptive to the basal lipid membrane voltage.^21^ Hence, a potential limitation of Cre-dependent expression of ChR2 is that we may not be able to tune and distribute expression of ChR2. However, in this study, we used a doxycycline-inducible and skeletal muscle-specific Cre which improved the specificity and can allow for temporal control of ChR2 expression for future studies.

Future studies should focus on whether ChR2 stimulation in skeletal muscle increases the leak currents within neighboring cell bodies (i.e. resting membrane potential, membrane resistance, and action potential in surrounding skeletal muscle cell-not exposed to light-changes compared to non-excited controls). This model is easily adjustable and, therefore, the stimulation protocol can be further optimized to reduce inter-pulse relaxations and improve the contractile profile under optical stimulation. In this study we showed the feasibility of optogenetic stimulation in Acta1-Cre Ai32 mice and to sustain muscle contraction for extended supraphysiological levels. It has been suggested that light stimulation might be more energy efficient in contraction of cardiomyocytes compared to electrical stimulation.^7^ At short pulses, AP morphology is similar between electrical and optogenetic stimulation; longer stimulation durations, however, result in apparent longer plateau phase that may be in part caused by the intrinsic negative feedback control of ChR2 activation.^19^ In future, effects of sustained prolonged contraction on excitation contraction coupling during light stimulation compared to electrical stimulation should be characterized. Lastly, measurement of *in vivo* electrophysiology in the light-sensitive cells under blue light using patch clamp analysis, and quantification of the delay in excitation in *in vivo* optogenetic stimulation compared to electrical stimulation would be informative in characterization of our proposed model.

Direct optogenetic stimulation shows promising application for restoration of function and atrophy caused by muscle denervation. Optogenetic muscle activation has recently been explored skeletal and facial muscle complexes.^11,12,22–24^ In mice, studies have shown direct optogenetic stimulation to improve functional metrics in denervated mouse whisker pads.^23^ Others have used Sim1-Ai32 transgenic mouse line, with non-specific expression in skeletal muscle t-tubule and sarcolemma, to attenuate denervation-caused skeletal muscle atrophy using repeated optogenetic stimulation.^12^ In addition to transgenic mouse models, transplantation of murine embryonic stem-cell derived motor neurons for targeted light stimulation of ChR2 in denervated sciatic nerve branch improved function of denervated muscle.^24^ However, applicability of optogenetic stimulation can be expanded to study load-dependent development and maturation of musculoskeletal tissues in juvenile mice. Muscle loading is critical for the healthy development of musculoskeletal tissues.^25–27^ In in case of enthesis, others have shown that unloading of this tissue using intermuscular injection of botulinum toxin A in supraspinatus muscle is disruptive to the development of functionally graded supraspinatus enthesis.^28^ Role of increased muscle loading on structure and function of the developing enthesis, however, is not well understood in the field due technical limitations related to stimulating the growing muscle in neonatal mice using standard approaches such as electrical stimulation and treadmill training. Direct optogenetic stimulation can be widely utilized as a tool for non-invasive tools induce muscle contraction for better understanding the mechanobiological cascades involved in formation and homeostasis of the enthesis *in vivo*.

## Conclusion

*In vivo* and direct optogenetic stimulation of skeletal muscle is a promising tool that can also be used in conjunction with denervation models to study muscle regeneration. Optogenetics can be used as a non-invasive toolbox for investigating the complex mechanobiological cascades involved in development of skeletal tissues and effect of muscle loading on adjacent tissue (e.g., tendon) formation and adaptation. Our model has the potential to overcome the systemic challenges and limitations of the existing models for muscle loading in the study of development of tendon, enthesis, and bone.

## Acknowledgements

Thanks to Dr. Gwen Talham and Frank Warren for assistance with animal care; Matthew Hudson and Brittany Wilson for sharing use of the Aurora 3-in-1 *in vivo* system; Jaclyn Soulas for assistance with whole mount imaging of the triceps surae; and Harrah Newman for preliminary power meter studies.

