## Supplemental data for "Optogenetic activation of muscle contraction *in vivo*"

**Supplementary Figures**

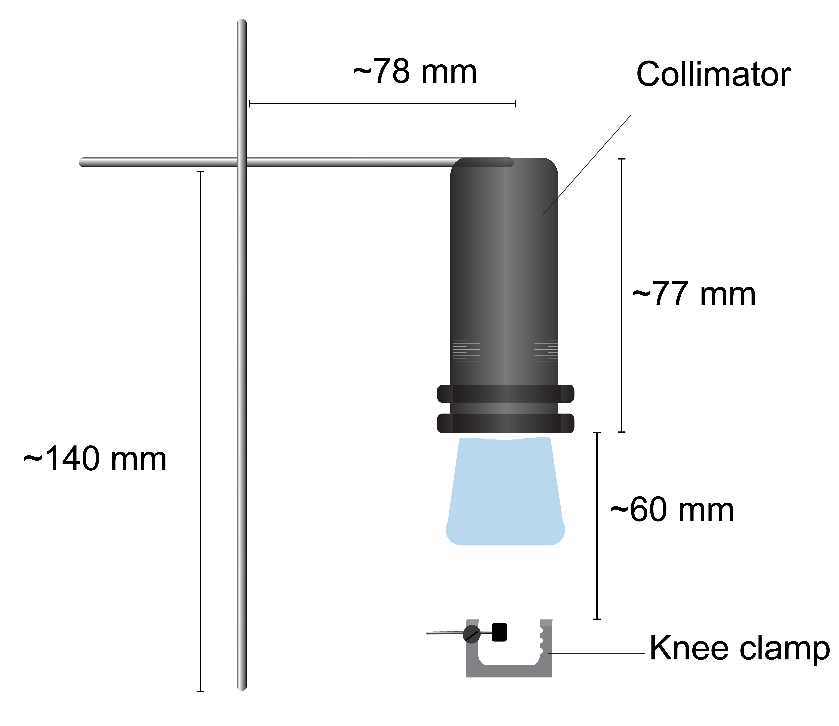

**Supplemental Figure 1.** Schematics showing experimental setup for optogenetic stimulation.

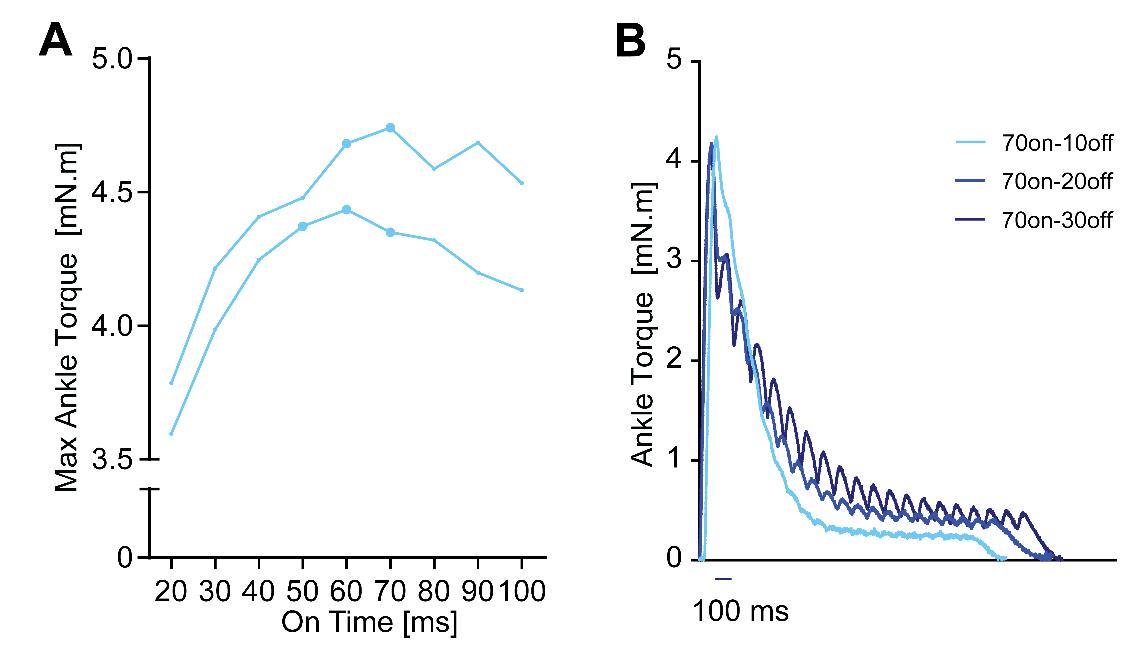

**Supplemental Figure 2.** Representative images showing contractile profile during (A) On and (B) Off time optimization. (A) Peak ankle torque occurs at 60-70ms On-time and (B) increase in the Off-time duration increases the sustain contractile ankle torque under pulsatile stimulation.

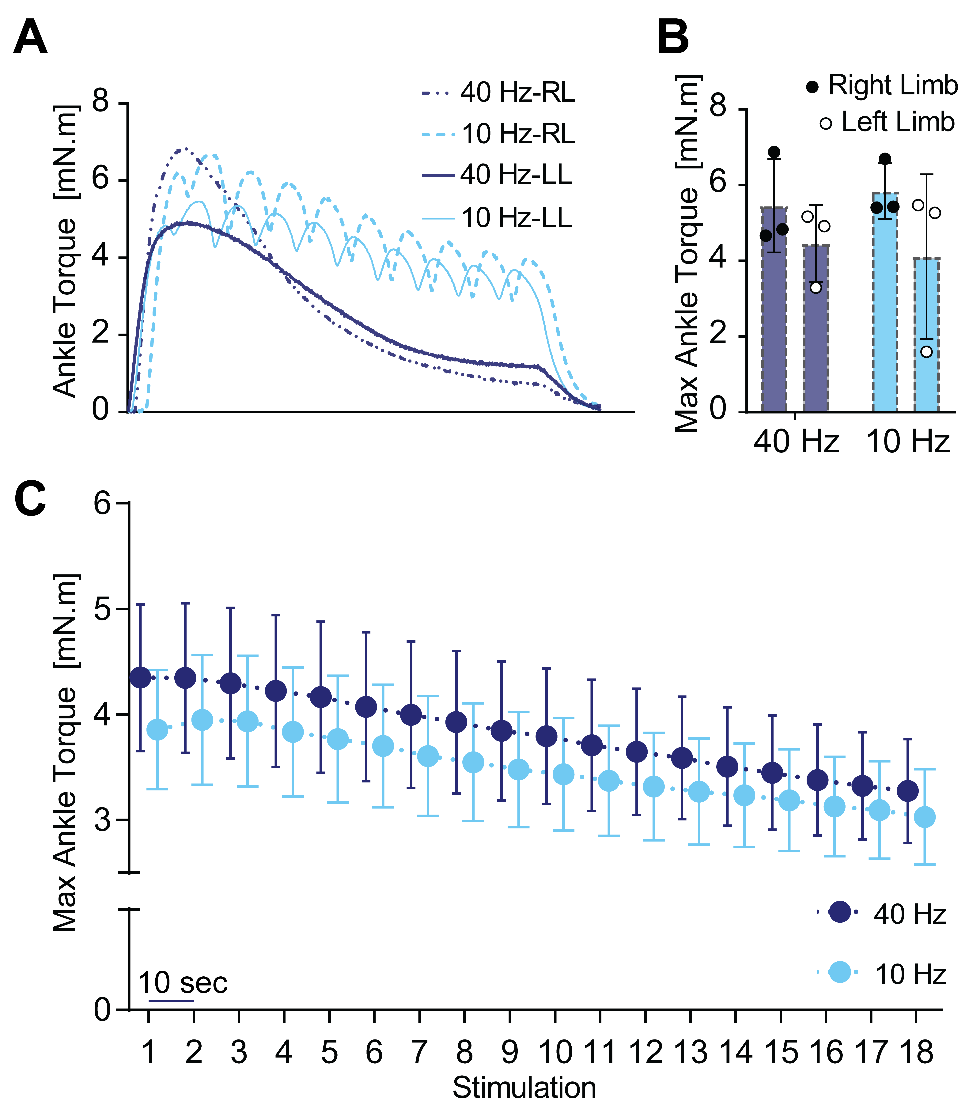

**Supplemental Figure 3.** Optogenetics can induce comparable contraction in contralateral limbs as shown by (A) representative contractile profile of contralateral limbs and (B) maximum contraction induced by under 10Hz and 40Hz optogenetic stimulation. (C) Decay in peak ankle torque after repeated 1-second duration optogenetic stimulation “bouts” was comparable between 10Hz and 40Hz with 10 seconds rest between stimulations. Total duration of repeated stimulations was 3 minutes including rest.

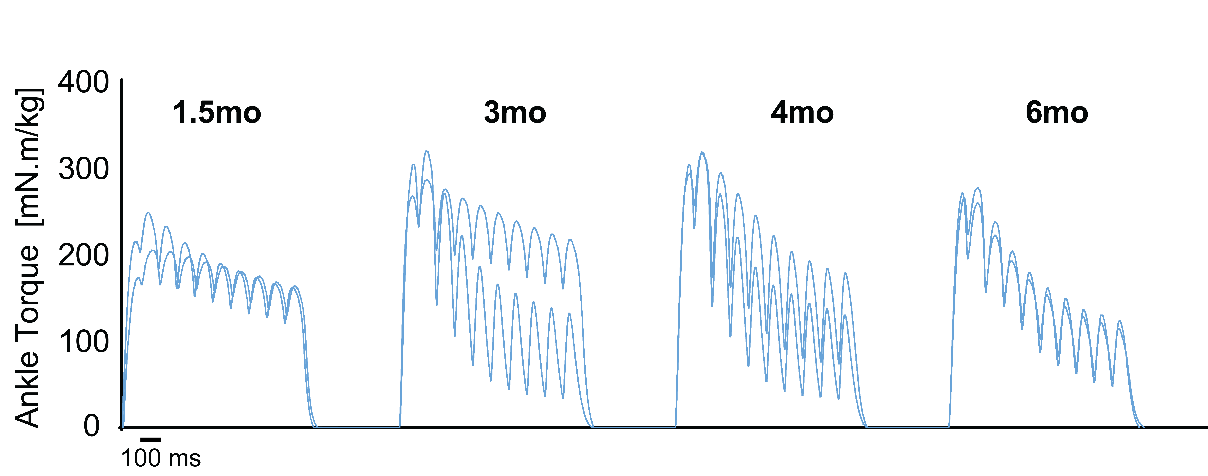

**Supplemental Figure 4.** Optogenetics stimulation of TS muscle can induce comparable results in young adult mice of 1.5 to 6 months of age.

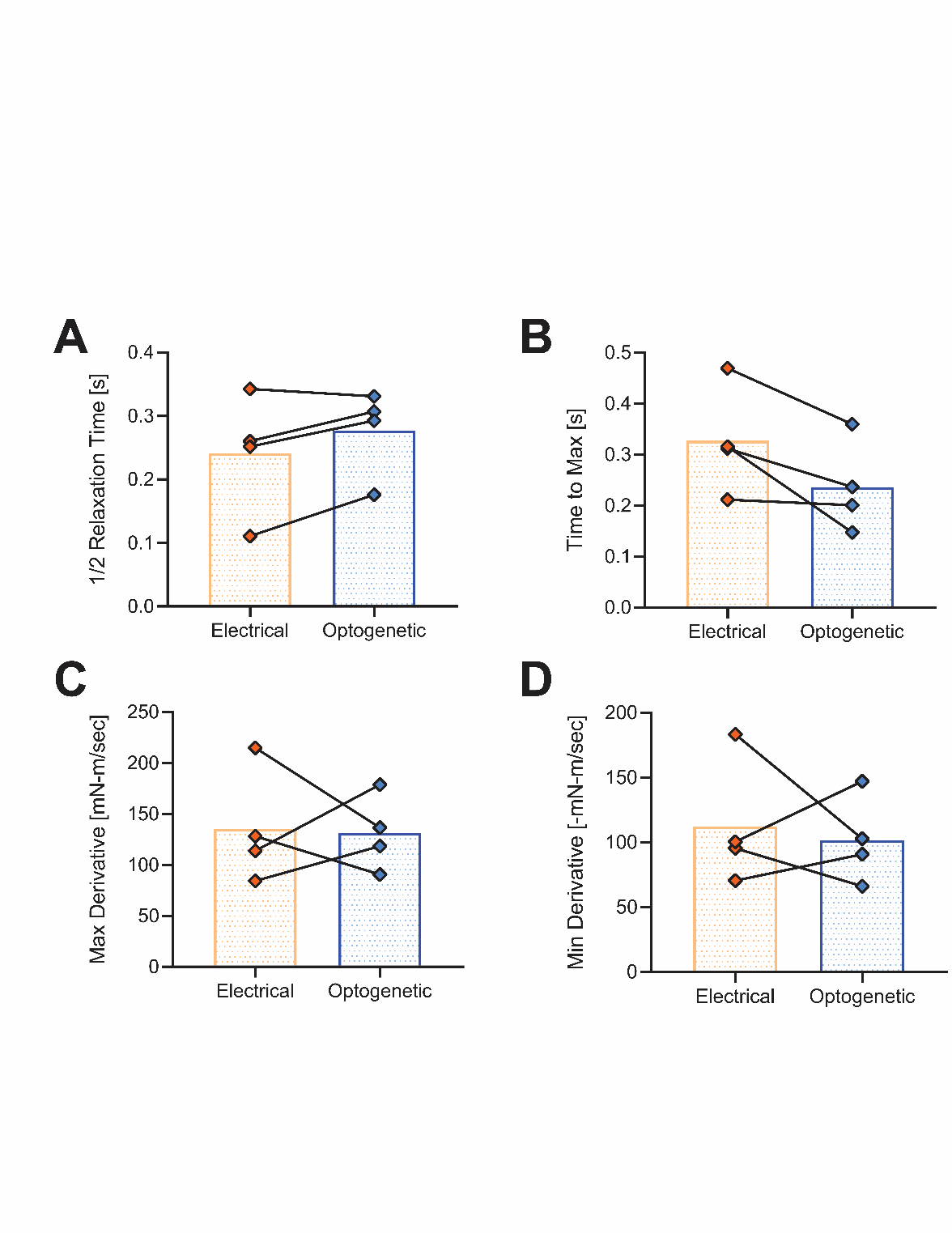

**Supplemental Figure 5.** Results from muscle analysis show that (A) half relaxation time, (B) time to maximum, (C) maximum derivative, and (D) minimum derivative are comparable between electrical and optogenetic stimulation.
